# A multiplex PCR assay for the detection of *Cryptosporidium* species and simultaneous differentiation of *Cryptosporidium hominis, Cryptosporidium parvum* in clinical stool samples

**DOI:** 10.1101/2023.03.22.533796

**Authors:** Manish Katiyar, Shashiraja Padukone, Reena Gulati, Rakesh Singh

## Abstract

*Cryptosporidium hominis* and *Cryptosporidium parvum* are responsible for more than 90% of the global cryptosporidiosis. Species identification is done by amplification of small subunit ribosomal ribonucleic acid (SSU rRNA) gene, followed by sequencing. We have developed a multiplex polymerase chain reaction (mPCR) assay which detect *Cryptosporidium* spp. and differentiates *C. hominis* and *C. parvum* from stool samples without the need of post amplification sequencing. Nine new set of primers for mPCR assay were designed and the mPCR assay was standardized with known positive *Cryptosporidium* DNA template. Best result with three sets of primers that amplifies 436 bp for all *Cryptosporidium* spp., 577 bp for *C. hominis* and 287 bp for *C. parvum*. In addition, thirty-five positive and thirty-five negative *Cryptosporidium* stool samples identified by the gold standard nested 18S rRNA PCR-sequencing assay were tested by mPCR. The sensitivity of the mPCR are 100%, 92.9%, and 87.5% for *Cryptosporidium* spp., *C. hominis*, and *C. parvum* respectively while specificity is 100% for all the three primers. No cross-reactivity was observed by the new mPCR assay when tested with five known DNA sample of *Cystoisospora belli* and two known DNA sample of *Cyclospora cayetanensis*, available in our laboratory from the clinical stool samples. A single species-specific mPCR product of *C. hominis* and *C. parvum* were sequenced and deposited in GenBank database with the accession no MT862538 and MT875168 respectively. The mPCR assay is developed which differentiates *C. hominis*, and *C. parvum* in a single test run of amplification and without the need for RFLP or sequencing. Although it less sensitive than 18S rRNA PCR-sequencing assay, but 100% specific, rapid, cost-effective and suitable for making diagnosis of cryptosporidiosis especially in developing countries.

**Highlights:**

- Novel mPCR assay can detect all *Cryptosporidium* species
- The sensitivity of the mPCR were 100%, 92.9%, and 87.5% for the primers designed to detect *Cryptosporidium* genus, *C. hominis* and *C. parvum* species respectively.
- No cross reactivity detected with newly developed mPCR assuring 100% specificity.
- The developed mPCR assay is a robust, specific, reproducible, rapid and cost-effective molecular assay for the diagnosis of cryptosporidiosis.
- Assay is useful in molecular diagnosis of cryptosporidiosis, especially in developing countries.

## 1. INTRODUCTION

*Cryptosporidium* species infect a variety of hosts which include humans and animals. It is one of the most common causes of global death from diarrhea in children under five years (Checkley et al., 2015; Vanathy et al., 2017). *Cryptosporidium hominis* and *Cryptosporidium parvum* accounts for almost 90% of human cryptosporidiosis (Desai et al., 2012; Guo et al., 2015; Liu et al., 2020; Wang et al., 2018). Vrious *Cryptosporidium* spp. which cause human cryptosporidiosis are *Cryptosporidium meleagridis, Cryptosporidium andersoni, Cryptosporidium ubiquitum, Cryptosporidium muris, Cryptosporidium baileyi, Cryptosporidium felis, Cryptosporidium canis, Cryptosporidium suis, Cryptosporidium cuniculus* and *Cryptosporidium* chipmunk genotype I. In India, *C. hominis* and *C. parvum* are responsible for almost all the outbreaks of cryptosporidiosis (Vanathy et al., 2017). *C. hominis* are common in India, Brazil, Pakistan, and Peru, while infections by *C. hominis* and *C. parvum* are almost equally distributed in European nations, Australia and the USA (Peralta et al., 2016). *C. parvum* have a broad host range, infect domesticated animals, especially cattle and humans acquire it through zoonotic transmission whereas *C. hominis* is acquired through anthroponotic transmission (Xiao, 2010). Evolving new diagnostic techniques for species and subtype identification is essential for diagnotics, understanding the pattern of transmission, virulence, monitoring prevalence, epidemiology of outbreaks, and development of appropriate control programs.

Microscopic identification of oocysts of *Cryptosporidium* spp. in stool smears is the most common method and is done by examining stool smears stained either with modified ziehl-neelsen method or auramine O method. Multiple stool samples over three days may be tested for diagnosis due to intermitted shedding of oocysts (Murphy et al., 2011; Vanathy et al., 2017). Diagnostic reliability also depends on the technical skill and expertise of the microscopist. Immunofluorescence microscopy is more sensitive but expensive for routine diagnostic use in developing countries (Checkley et al., 2015). Diagnostic sensitivity of commercially available enzyme-linked immunosorbent assays (ELISA) or immuno-chromatographic (ICT) methods is highly variable (Checkley et al., 2015), besides none of these methods differentiate the species of *Cryptosporidium* (Thompson et al., 1994).

Polymerase chain reaction (PCR) is now considered as a gold standard test for the diagnosis of cryptosporidiosis. Amplification of conserved gene of *Cryptosporidium* spp. i.e small subunit ribosomal ribonucleic acid (SSU rRNA) gene, followed by either Restriction Fragment Length Polymorphism (RFLP) or sequencing is a widely used molecular technique (Xiao, 2010)[8]. Other gene targets such as TRAP C1, actin, GP60, internal transcribed spacers (ITS), COWP, HSP 70, and DHFR have also been used with varying sensitivities (Fayer et al., 2000; Gibbons et al., 1998; Morgan et al., 1999; Ng et al., 2006; Pedraza-Díaz et al., 2001; Spano et al., 1998). Real-time PCR with the fluorescent probe-based method is highly sensitive and specific technique, but high cost makes its widespread implementation impractical in developing countries (Murphy et al., 2011; Staggs et al., 2013). Therefore a rapid, cost-effective, sensitive and specific laboratory technique is the current need in many developing countries where cryptosporidiosis is prevalent. With this background, we studied the target genes specific for *C. hominis* and *C. parvum* and combined them with genus-specific 18S rRNA gene to develop a multiplex PCR (mPCR).

## 2. MATERIALS AND METHODS

### 2.1. Ethical statement

The study was conducted at a teriary care teaching hospital and it was approved by the Institutional Ethics Committee (Human studies).

### 2.2. Positive control

Deoxyribonucleic acid (DNA) positive control of *C. hominis, C. parvum, C. meleagridis, C. felis, C. andersoni* were obtained from Christian Medical College (CMC) Vellore, India and were used as a positive control for standardization of PCR and mPCR. positive control were stored at -20^0^C until use.

### 2.3. Clinical stool sample

Seventy clinical stool samples which were tested by the molecular goal standard nested 18S rRNA PCR assay were available in our diagnostic laboratory. Thirty-five stool samples were positive for *Cryptosporidium* spp. and 35 were negative for *Cryptosporidium* spp (unpublished data). DNA sample were extracted from stool samples using QIAmp DNA Stool Mini Kit (Qiagen, Hilden, Germany) as per the manufacturer’s instructions. The quality and quantity of the extracted DNA sample were evaluated with NanoDrop 2000C (ThermoFisher, Massachusetts, USA) and stored at –20^0^C until use. Nested 18S rRNA PCR assay was performed by the method as described by Xiao et al. with minor modification. Thirty-five stool samples yielded positive products of 850 bp in 1.2% gel electrophoresis (Xiao et al., 1999). Species of *Cryptosporidium* isolates were identified by sequencing the amplicon products with ABI 3730xl sequencer after purification by FavorPrep GEL/PCR purification mini kit (Cat. No: FAGCK001-1, Ping-Tung, Taiwan) as per manufacturer’s instructions. Sequenced data were visualized using FinchTV (v1.4.0, Geospiza Inc, Seattle, USA). DNA Baser Assembler (v5.15.0, Heracle BioSoft, Arges, Romania) and obtained consensus sequences. Consolidated sequences were searched in GenBank by BLASTn for the best match. Thirty-five positive isolates were identified as sixteen *C. parvum*, fourteen *C. hominis*, two each of *C. felis* and *C. canis* and one *C. muris*. These were used to validate the mPCR. The sequence data have been deposited in the GenBank database under accession no. MW116641 to MW116659, MT835257 to MT835271, and MW228121.

### 2.4. Primer designing of mPCR

A Basic Local Alignment Search Tool (BLAST http://www.ncbi.nlm.nih.gov/blast/Blast.cgi) was used to search for the conserved sequences of *Cryptosporidium* rRNA gene. *C. parvum* MT 18S rRNA (Accession number AF161856.1) was searched as a query using NCBI-BLASTn with default settings. The resulting sequences with 100% query coverage and > 95% identity were retrieved and aligned with ClustalW in MEGA v7 by multiple sequence alignment to identify common regions suitable for genus-specific primer (Kumar et al., 2016; Thompson et al., 1994). Primers were designed based on regions that are conserved using Primer-3 (http://www.bioinformatics.nl/cgi-bin/primer3plus/primer3plus.cgi). Chro.50011, the only gene specific for *C. hominis* and, cgd6.5510, cgd8_680, cgd8_690, cgd6_5490, cgd5_4580, and cgd5_4610 genes unique for *C. parvum* were used to design specific primers for *C. hominis* and *C. parvum* respectively (Guo et al., 2015). Primers were also examined in silico in SnapGene software (v1.1.3, Chicago, USA). The three sets of primers – *Cryptosporidium* spp., *C. hominis*, and *C. parvum* were multiplexed and were tested for the best possible amplification product.

### 2.5. Standardization of PCR

Uniplex PCRs were initially tested by using all the designed primers using *Cryptosporidium* positive controls and nuclease-free water as negative control. Gradient PCR was performed to standardize the annealing temperature. Cycling conditions for each PCR were standardized for initial thermal denaturation, annealing temperature, extension temperatures, number of cycles, and primer concentration. All amplifications were performed with commercial 2 X Taq DNA Polymerase master mixes (Amplicon, Odense, Denmark) in a final volume of 25 μl on Agilent SureCycler 8800 (Agilent Technologies, California, USA). Electrophoresis was conducted with 1.2% agarose gel and visualized in the Biorad gel documentation system. PCR amplicons that showed the expected product size and matched with product size of the positive control were considered positive, and the rest were considered negative. Multiplexing was done by using three sets of primers i.e. primers for *Cryptosporidium* spp. and primers specific for *C. hominis* and *C. parvum*. Positive control of *C. parvum* and *C. hominis* was mixed in equal quantities, two such mixes were constituted and tested by mPCR to analyse its ability to detect different *Cryptosporidium* species in a mixed sample.

### 2.6. Sensitivity and specificity of mPCR

Sensitivity and specificity of the mPCR was determined by considering nested 18S rRNA PCR as a gold standard reference molecular assay. Representative amplicons of mPCR product were also sequenced to determine the specificity of the mPCR.

### 2.7. Test for cross-reactivity

Cross-reactivity of the mPCR assay were checked by testing mPCR with known five DNA sample of *Cystoisospora belli* and two DNA sample of *Cyclospora cayetanensis*. These DNA controls were detected and available in our laboratory.

### 2.8. Evaluation of diagnostic reproducibility

Diagnostic reproducibility and repeatability of the mPCR assay were evaluated by testing on two other PCR platforms Applied Biosystems (Veriti Thermal Cycler, Catalogue No. 4375786, Waltham, USA) and Eppendorf (master cycle nexus, Catalogue No. 6331000017, Hamburg, Germany). Five randomly selected positive samples each of *C. parvum* and *C. hominis* were tested on both platforms. PCR reproducibility were also tested with PCR master mix obtained from other manufacturers, i.e. Takara (EmeraldAmp® GT PCR Master Mix, RR310B, California, USA), Promega (GoTaq® Master Mix, Catalogue No. M7122, Madison, USA), and Qiagen (Taq PCR Master Mix Kit, Catalogue No. 201443, Hilden, Germany) using Agilent SureCycler PCR platform.

## 3. RESULTS

### 3.1 Primer designing

A total of nine sets of primers were designed – one for *Cryptosporidium* spp., two sets specific for *C. hominis* and 6 sets for *C. parvum* (Table 1). They were examined in silico in SnapGene software (v1.1.3) and found to be satisfactory (Figure 1).

**Table 1.**
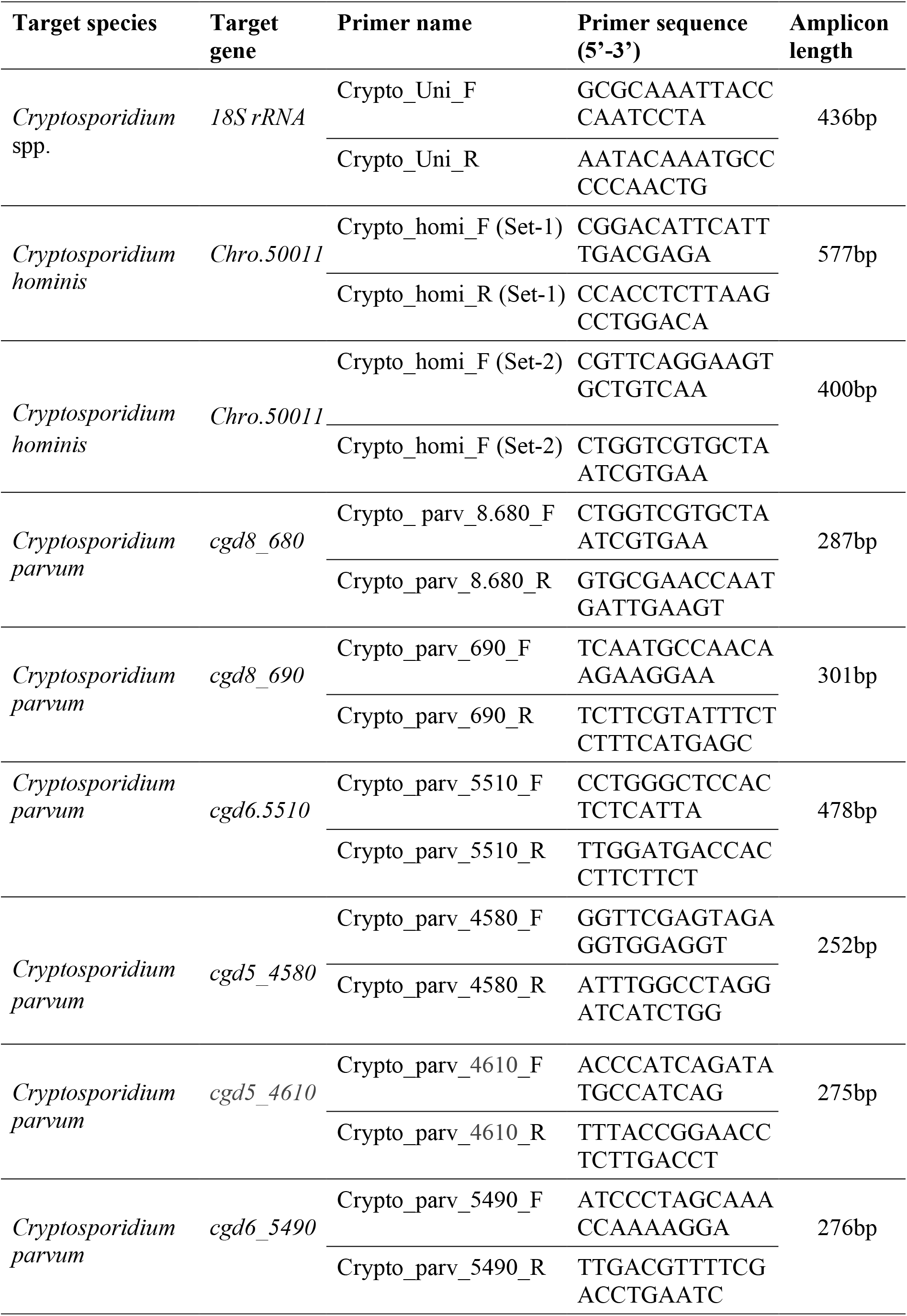
List of primers designed for *Cryptosporidium* spp.

**Figure 1.**
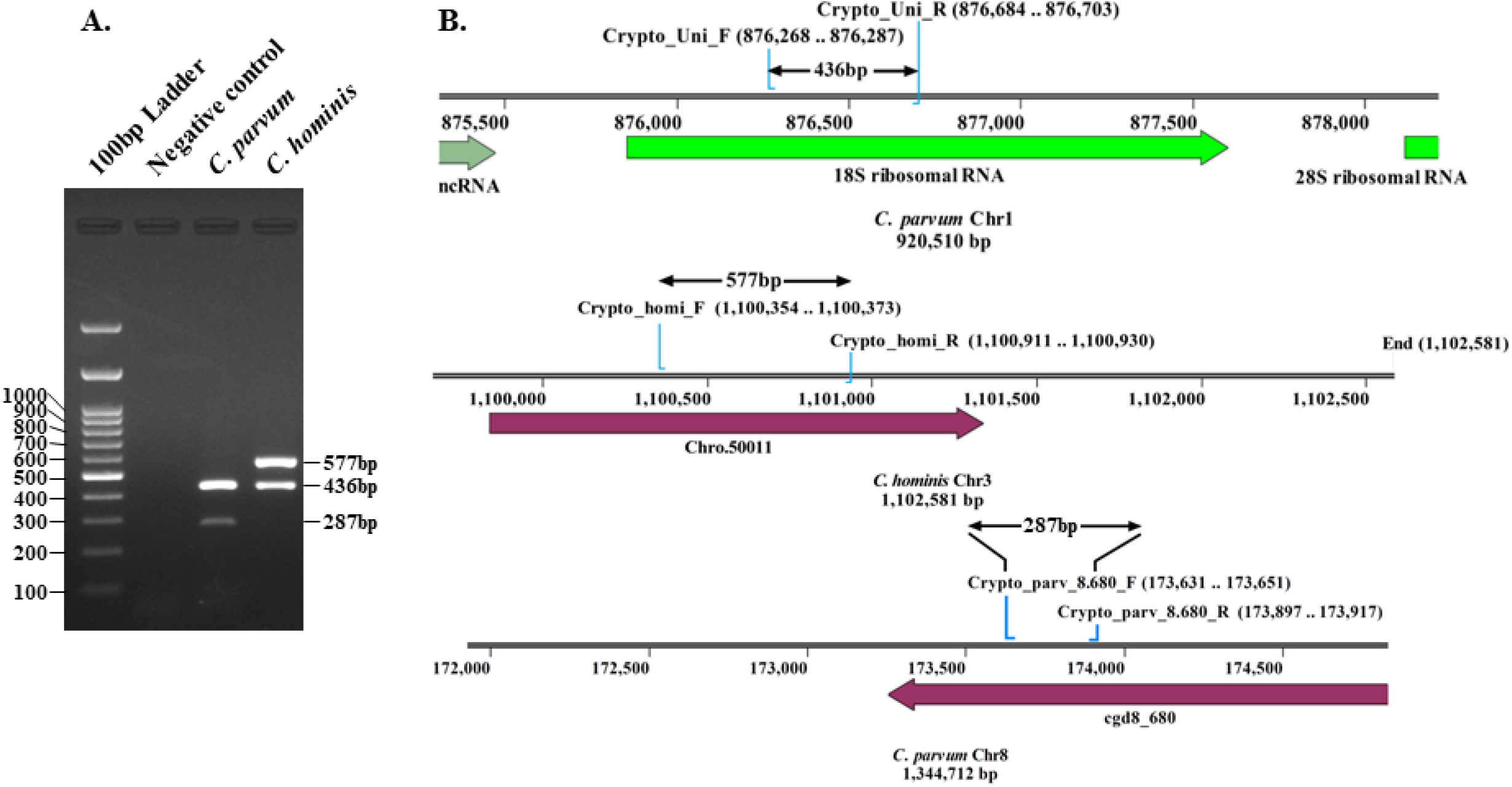
(A) Gel electrophoresis of mPCR with positive *Cryptosporidium* DNA material; Lane 1: 100 bp DNA ladder; Lane 2: Negative control; Lane 3: 287 bp and 436 bp indicative of *C. parvum*; Lane 4: 436 bp and 577 bp indicative of *C. hominis* (B) Primer-binding sites and amplicon lengths of the three primers of mPCR by SnapGene software (v1.1.3)

### 3.2. Standardization of mPCR

Uniplex amplification of *Cryptosporidium* 18S rRNA (436bp) gene, species-specific, Chro.50011 gene by set-1 primer (577bp), Chro.50011 gene by set-2 primer (400bp), cgd8_680 (287 bp), cgd8_690 (301 bp), cgd6.5510 (478 bp), cgd5_4580 (252 bp), cgd5_4610 (275 bp) and cgd6_5490 (276 bp) genes were performed successfully with positive control of *Cryptosporidium* spp. Multiplexing was done by using three sets of primers, and of the six primers for genes (cgd8_680, cgd8_690, cgd6_5510, cgd6_5490, cgd5_4580, and cgd5_4610) targeted for *C. parvum* species-specific amplification, primers for cgd8_680 gene produced the best results when multiplexed with *C. hominis* Chro.50011 set-1 primers. The best panel of the primers for the mPCR assay were 436 bp for 18S rRNA gene for *Cryptosporidium* spp., primer for 577 bp for Chro.50011 gene specific for *C. hominis*, and 287 bp for cgd8_680 gene specific for *C. parvum* (Figure 1). The mPCR showed expected positive results with all the positive controls of *Cryptosporium* spp. Which include *C. hominis, C. parvum, C. meleagridis, C. andersoni*, and *C. felis*.

The optimal PCR cycling conditions of mPCR were: initial denaturation at 94^0^C for 5 minutes followed by 35 cycles of 94^0^C for 45 seconds, 60^0^C for 45 seconds, and 72^0^C for 45 seconds, followed by a final extension at 72^0^C for 5 minutes. PCR reaction volume was standardized as 12.5 μl of commercial 2X Taq DNA polymerase master mix, 2 μl of DNA template, 0.3 μM each of *Cryptosporidium* spp. and *C. hominis* primers, 0.5 μM of *C. parvum* primers, and rest nuclease-free water in a final volume of 25 μl.

### 3.3. Sensitivity and Specificity of mPCR

The performance of mPCR was validated with gold standard nested 18S rRNA PCR. It gave a concordant result for all 70 samples tested - 35 positive and 35 negative samples (Figure 2 Lane 4 to 6). The sensitivity and specificity of mPCR are 100% for genus-specific primers of 18S rRNA genes.

**Figure 2.**
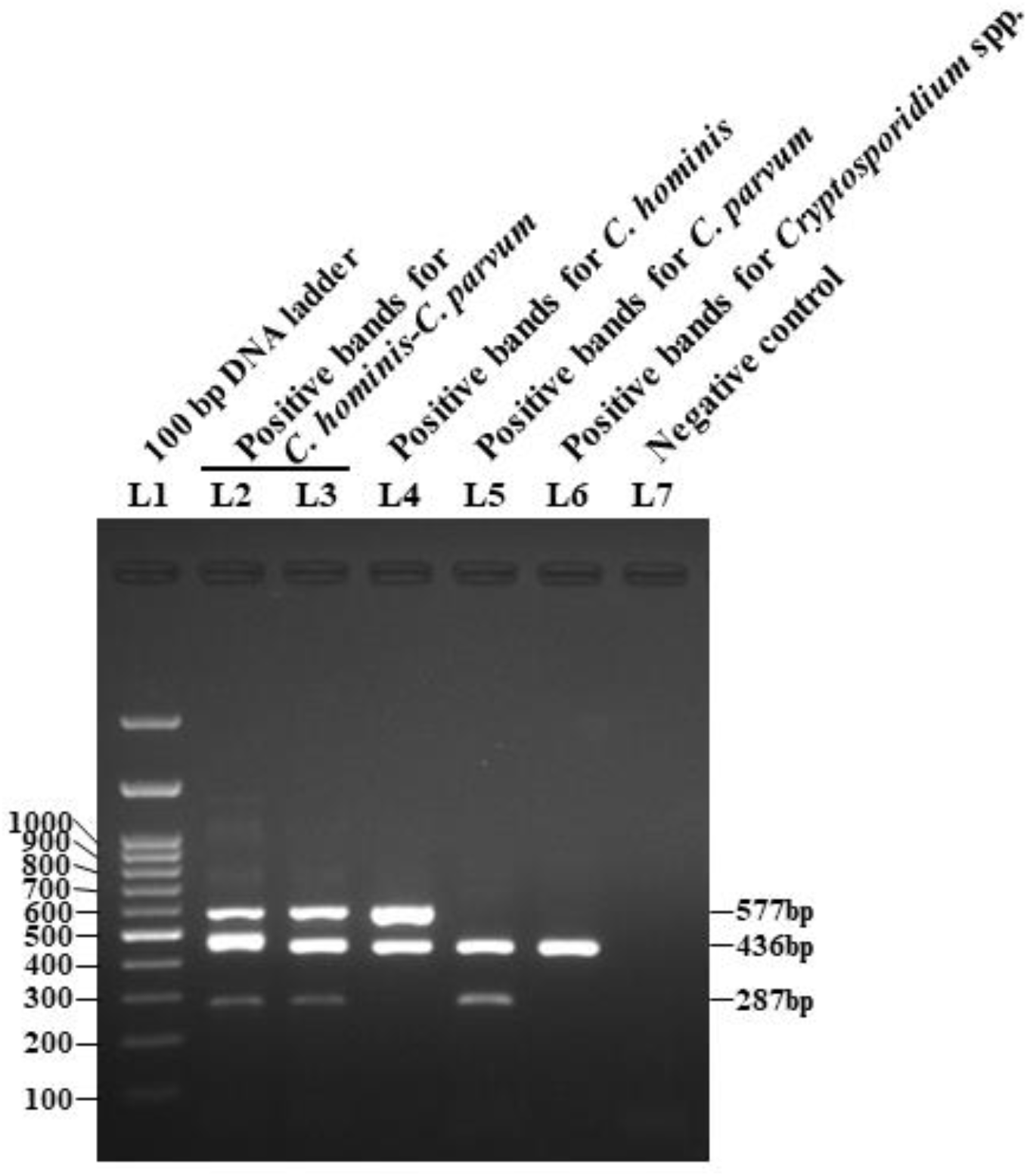
Gel electrophoresis of mPCR assay for the detection of *C. hominis, C. parvum* and NHNPC; Lane 1: 100 bp DNA ladder; Lane 2 & 3: 287 bp, 436 bp and 577 bp bands indicative of mixture of *C. hominis* and *C. parvum*; Lane 4: 436 bp and 577 bp bands indicative of *C. hominis*; Lane 5: 287 bp and 436 bp bands indicative of *C. parvum*; Lane 6: Single positive band indicative of NHNPC; Lane 7: Negative control

The mPCR was able to identify 13 of 14 *C. hominis* positive stool samples i.e sensitivity of 92.9% for *C. hominis* and detected 14 out of 16 *C. parvum* positive stool samples i.e sensitivity of 87.5% for *C. parvum*. It did not detect any false positive result and therefore it is 100% specific for *C. hominis* and *C. parvum*.

Representative amplicons of mPCR product were purified, sequenced, and deposited in the GenBank database with the accession no MT796062 for *C. hominis*, MT796063 for *C. parvum*, and MT796064 for *C. felis*, which were amplified by using genus-specific *Cryptosporidium* spp. primers. Similarly, representative amplicon of mPCR product for Chro.50011 gene specific for *C. hominis* and cgd8_680 gene product specific for *C. parvum* were sequenced and deposited in GenBank database with the accession no MT862538 and MT875168 respectively.

DNA extract material of *C. hominis* and *C. parvum* were mixed in equal quantities, and two such mixes were constituted. The mPCR accurately detected mixed infection in both the constituted DNA extract samples (Figure 2 Lane 2 & 3).

### 3.4. Test for cross-reactivity

No cross-reactivity was observed by the new mPCR assay when tested with five DNA extract material of *C. belli* and two DNA extract material of *C. cayetanensis*, confirming 100% specificity of the method.

### 3.5. Evaluation of reproducibility

Diagnostic reproducibility and repeatability of the PCR assay were evaluated by testing on two other PCR platforms - Applied Biosystems and Eppendorf; and PCR master mix obtained from other manufactures – Takara, Promega and Qiagen. A positive band was observed in all five random positive samples tested irrespective of the PCR platform and the PCR reagents.

## 4. DISCUSSION

Cryptosporidiosis in humans is predominantly caused by *C. hominis* and *C. parvum*. In some occasion human cryptosporidiosis is caused by species other than *C. hominis* and *C. parvum*. Species identification of *Cryptosporidium* is not routinely performed due to the lack of robust, reliable, and rapid diagnostic method. The prevalence of individual species causing cryptosporidiosis may be useful for implementation of appropriate control programs. Species identification is reliably done by molecular assays. Hence, molecular diagnostic techniques are the present research subject (Bouzid et al., 2016; Santin and Zarlenga et al., 2009; Shrivastava et al., 2020). In this context, we developed a sensitive, reliable, cost-effective and specific multiplex PCR technique to identify the two common prevalent *Cryptosporidium* spp. in a single test run of amplification and without the need for RFLP or sequencing. For *C. hominis*, the only known specific gene Chro.50011 is located on chromosome 3, encodes a hypothetical protein of 489 amino acids (Guo et al., 2015; Shrivastava et al., 2020). Chro.50011 gene is present at a telomeric location, are involved in host specificity and virulence (Guo et al., 2015; Nader et al., 2019). There are several genes, such as cgd8_680, cgd8_690, cgd6_5510, cgd6_5490, cgd5_4580, and cgd5_4610, that are unique for *C. parvum* (Guo et al., 2015). The products of these polymorphic genes belong to a new family of telomerically-encoded *Cryptosporidium* proteins. The cgd8_680 gene is located on chromosome 8 of *C. parvum*. Of the six *C. parvum*-specific gene targets, cgd8_680, produced the best results in mPCR.

The best panel of the primers for the mPCR assay identified here are 436 bp for 18S rRNA gene for *Cryptosporidium* spp., primer for 577 bp for Chro.50011 gene specific for *C. hominis*, and 287 bp for cgd8_680 gene specific for *C. parvum*. This mPCR identifies the genus level and species-specific distinct fragments of different product sizes, enabling differentiation between *C. hominis* and *C. parvum* in a single test run of amplification. Shrivastava et al., have recently developed a real time PCR for the detection of *C. hominis* / *C. parvum* (Shrivastava et al., 2020). In their method, the species identification was not done, whereas our mPCR detects *C. hominis, C. parvum* and NHNPC. Sant et al., developed a multiplex PCR for detecting *C. parvum, Cryptosporidium bovis, Cryptosporidium ryanae*, and *Cryptosporidium andersoni* in infected cattle. It was developed for detecting cryptosporidiosis in cattle and therefore *C. hominis* was not included in their molecular assay (Santin and Zarlenga et al., 2009).

Guo et al., stated that cgd8_680 gene is specific to *C. parvum* (Guo et al., 2015). but recently Xu et al. reported that the cgd8_680 gene of *C. parvum* has paralogs in *C. hominis* (Chro.80081), *C. meleagridis* (C_mele_47675.3516), *C. ubiqitum* (cubi03292) and *C*.*baileyi* (C_bai_692.2837) (Xu et al., 2019). However, in our study, we did not find any amplification of the *C. hominis Chro*.*80011* gene by cgd8_680 primers of *C. parvum* among 14 known isolates of *C. hominis*. Xu et al. also observed that *Chro*.*50011* gene is specific to *C. hominis* (Xu et al., 2019).

A single species-specific mPCR product for *C. hominis* and *C. parvum* were sequenced and deposited in GenBank database with the accession no MT862538 and MT875168 respectively. BLASTn search of MT862538 and MT875168 yielded only two more sequences in GenBank database for each search. It indicates lack of research in this part of *Cryptosporidium* genome. BLASTn search of MT862538 matches 100% sequence data of *C. hominis* with accession number XM_660592 published on 1^st^ November, 2008. The other sequence data is of Aquila chrysaetos chrysaetos genome, matches with 100% identity and 5% query coverage which is completely unrelated to *Cryptosporidium* spp. testing on human stool samples and hence does not affect the test result. On the other hand MT875168 matches with 94.42% identity and 95% query coverage with the only two sequence data of *C. parvum* available in GenBank with accession number CP044415 published on 5^st^ November, 2020 and XM_625527 published on 1^st^ November, 2008. BLASTn search of the species-specific mPCR product for *C. hominis* and *C. parvum* indicate that the mPCR assay is specific, noval and not much studied earlier.

There have been recent reports of development of real-time PCR for the detection of *Cryptosporidium* spp. as well as the simultaneous identification of *C. hominis* and *C. parvum* (Bouzid et al., 2016; Hadfield et al., 2011; Mary et al., 2013). It is used at a reference center in a developed country. Conventional PCR assays are cost-effective than real time PCR assays. This current mPCR assay is a single test run of PCR amplification assay and is cost effective than the current gold standard nested PCR assay which requires two runs of PCR amplification. The cost of the current mPCR is further reduced because it does not require RFLP or sequencing whereas gold standard test needs either RFLP or sequencing for species identification. The sensitivity of the mPCR are 100%, 92.9%, and 87.5% for genus-specific primers of 18S rRNA genes, primers for species-specific Chro.50011 gene for *C. hominis*, and primers for species-specific cgd8_680 gene for *C. parvum*, respectively compared to the gold standard nested 18S rRNA PCR–sequencing assay. The mPCR takes advantage of the multi copy number of 18S rRNA genes of *Cryptosporidium* spp. It is used for genus level detection and therefore makes the test 100% sensitive. The lower sensitivity of *C. parvum* in the current mPCR may be due to two reasons. First, the AT content of the cgd8_680 gene is only 50%, much lower than the genome average of ∼70%, which makes it difficult to design optimal primers for PCR. Secondly, the mPCR was performed on the stocked DNA material in the laboratory and we have an opinion that the sensitivity of current mPCR would be higher if tested on freshly collected stool sample. However, the specificity of the mPCR is 100% for all three sets of primers.

The mPCR assay has been validated through *in-silico* examination, expected product size in gel electrophoresis, and sequencing of representative genus-specific and species-specific PCR products. It does not amplify other intestinal coccidian parasitic DNA when tested with *C. belli* and *C. cayetanensis*. It was also validated on different PCR platforms and with consumables from other manufacturers, confirming its reproducibility, making it technically less demanding and easily introduced in any molecular laboratory.

We are having opinon that our mPCR assay can be useful in place of gold standard 18S rRNA nested PCR-RFLP/sequencing assay in many countries where *C. hominis* and *C. parvum* are the common agents of cryptosporidiosis. The current mPCR can also detect mixed infections of *C. hominis* and *C. parvum* in stool samples. Our mPCR assay can be further developed into a commercial cost-effective diagnostic kit. Besides its epidemiological applicability, this assay can play a significant role in the diagnosis of a patient with diarrhea, especially in developing countries.

## 5. CONCLUSIONS

The developed mPCR assay is a robust, specific, reproducible, rapid and cost effective molecular assay that identifies *C. hominis, C. parvum*, NHNPC and mixture of *C. hominis* and *C. parvum*. in a single test run of amplification from a stool sample. Although it is less sensitive than gold standard 18S rRNA nested PCR-RFLP/sequencing assay, it is still useful in molecular diagnosis of cryptosporidiosis, especially in developing countries.

## REFERENCES

Bouzid, M., Elwin, K., Nader, J.L., Chalmers, R.M., Hunter, P.R., Tyler, K.M., 2016. Novel real-time PCR assays for the specific detection of human infective Cryptosporidium species. Virulence 7, 395–399. https://doi.org/10.1080/21505594.2016.1149670.

Checkley, W., White Jr, A.C., Jaganath, D., Arrowood, M.J., Chalmers, R.M., Chen, X.M., Fayer, R., Griffiths, J.K., Guerrant, R.L., Hedstrom, L., Huston, C.D., Kotloff, K.L., Kang, G., Mead, J.R., Miller, M., Petri, W.A. Jr, Priest, J.W., Roos, D.S., Striepen, B., Thompson, R.C., Ward H.D., Van Voorhis, W.A., Xiao L., Zhu, G., Houpt, E.R., 2015. A review of the global burden, novel diagnostics, therapeutics, and vaccine targets for Cryptosporidium. Lancet Infect. Dis. 15, 85–94. https://doi.org/10.1016/S1473-3099(14)70772-8.

Desai, N., Sarkar, R., Kang, G., 2012. Cryptosporidiosis: an under-recognized public health problem. Trop. Parasitol. 2, 91–98. https://doi.org/10.4103/2229-5070.105173.

Fayer, R., Morgan, U., Upton, S.J., 2000. Epidemiology of Cryptosporidium: transmission, detection and identification. Int. J. Parasitol. 30, 1305–1322. https://doi.org/10.1016/S0020-7519(00)00135-1.

Gibbons, C.L., Gazzard, B.G., Ibrahim, M.A.A., Morris-Jones, S., Ong, C.S.L., Awad-El-Kariem, F.M., 1998. Correlation between markers of strain variation in Cryptosporidium parvum: evidence of clonality. Parasitol. Int. 47, 139–147. https://doi.org/10.1016/S1383-5769(98)00012-9.

Guo, Y., Tang, K., Rowe, L.A., Li, N., Roellig, D.M., Knipe, K., Frace, M., Yang, C., Feng, Y., Xiao, L., 2015. Comparative genomic analysis reveals occurrence of genetic recombination in virulent Cryptosporidium hominis subtypes and telomeric gene duplications in Cryptosporidium parvum. BMC Genomics 16, 320. https://doi.org/10.1186/s12864-015-1517-1.

Hadfield, S.J., Robinson, G., Elwin, K., Chalmers, R.M., 2011. Detection and differentiation of Cryptosporidium spp. in human clinical samples by use of real-time PCR. J. Clin. Microbiol. 49, 918–924. https://doi.org/10.1128/JCM.01733-10.

Kumar, S., Stecher, G., Tamura, K., 2016. MEGA7: molecular evolutionary genetics analysis version 7.0 for bigger datasets. Mol. Biol. Evol. 33, 1870–1874. https://doi:10.1093/molbev/msw054.

Liu, A., Gong, B., Liu, X., Shen, Y., Wu, Y., Zhang, W., Cao, J., 2020. A retrospective epidemiological analysis of human Cryptosporidium infection in China during the past three decades (1987-2018). PLoS Negl. Trop. Dis. 14, e0008146. https://doi.org/10.1371/journal.pntd.0008146.

Mary, C., Chapey, E., Dutoit, E., Guyot, K., Hasseine, L., Jeddi, F., Menotti, J., Paraud, C., Pomares, C., Rabodonirina, M., 2013. Multicentric evaluation of a new real-time PCR assay for quantification of Cryptosporidium spp. and identification of Cryptosporidium parvum and Cryptosporidium hominis. J. Clin. Microbiol. 51, 2556–2563. https://doi.org/10.1128/JCM.03458-12.

Morgan, U.M., Deplazes, P., Forbes, D.A., Spano, F., Hertzberg, H., Sargent, K.D., Elliot, A., Thompson, R.C.A., 1999. Sequence and PCR--RFLP analysis of the internal transcribed spacers of the rDNA repeat unit in isolates of Cryptosporidium from different hosts. Parasitology 118, 49–58. https://doi.org/10.1017/s0031182098003412.

Murphy, S.C., Hoogestraat, D.R., SenGupta, D.J., Prentice, J., Chakrapani, A., Cookson, B.T., 2011. Molecular diagnosis of cystoisosporiasis using extended-range PCR screening. J. Mol. Diagnostics 13, 359–362. https://doi.org/10.1016/j.jmoldx.2011.01.007.

Nader, J.L., Mathers, T.C., Ward, B.J., Pachebat, J.A., Swain, M.T., Robinson, G., Chalmers, R.M., Hunter, P.R., Van Oosterhout, C., Tyler, K.M., 2019. Evolutionary genomics of anthroponosis in Cryptosporidium. Nat. Microbiol. 4, 826–836. https://doi.org/10.1038/s41564-019-0377-x.

Ng, J., Pavlasek, I., Ryan, U., 2006. Identification of novel Cryptosporidium genotypes from avian hosts. Appl. Environ. Microbiol. 72, 7548–7553. https://doi.org/10.1128/AEM.01352-06

Pedraza-Díaz, S., Amar, C., Nichols, G.L., McLauchlin, J., 2001. Nested polymerase chain reaction for amplification of the Cryptosporidium oocyst wall protein gene. Emerg. Infect. Dis. 7, 49. https://doi.org/10.3201/eid0701.700049.

Peralta, R.H.S., Velásquez, J.N., Cunha, F. de S., Pantano, M.L., Sodré, F.C., Silva, S. da, Astudillo, O.G., Peralta, J.M., Carnevale, S., 2016. Genetic diversity of Cryptosporidium identified in clinical samples from cities in Brazil and Argentina. Mem. Inst. Oswaldo Cruz 111, 30–36. https://doi.org/10.1590/0074-02760150303.

Santin, M., Zarlenga, D.S., 2009. A multiplex polymerase chain reaction assay to simultaneously distinguish Cryptosporidium species of veterinary and public health concern in cattle. Vet. Parasitol. 166, 32–37. https://doi.org/10.1016/j.vetpar.2009.07.039.

Shrivastava, A.K., Panda, S., Kumar, S., Sahu, P.S., 2020. Two novel genomic DNA sequences as common diagnostic targets to detect Cryptosporidium hominis and Cryptosporidium parvum: Development of quantitative polymerase chain reaction assays, and clinical evaluation. Indian J. Med. Microbiol. 38, 430–440. https://doi.org/10.4103/ijmm.IJMM_20_114.

Spano, F., Putignani, L., Guida, S., Crisanti, A., 1998. Cryptosporidium parvum: PCR-RFLP analysis of the TRAP-C1 (thrombospondin-related adhesive protein of Cryptosporidium-1) gene discriminates between two alleles differentially associated with parasite isolates of animal and human origin. Exp. Parasitol. 90, 195–198. https://doi.org/10.1006/expr.1998.4324.

Staggs, S.E., Beckman, E.M., Keely, S.P., Mackwan, R., Ware, M.W., Moyer, A.P., Ferretti, J.A., Sayed, A., Xiao, L., Villegas, E.N., 2013. The applicability of TaqMan-based quantitative real-time PCR assays for detecting and enumerating Cryptosporidium spp. oocysts in the environment. PLoS One 8, e66562. https://doi.org/10.1371/journal.pone.0066562.

Thompson, J.D., Higgins, D.G., Gibson, T.J., 1994. ClustalW: improving the sensitivity of progressive multiple sequence alignment through sequence weighting, position-specific gap penalties and weight matrix choice. Nucleic Acids Res. 22, 4673–4680. https://doi.org/10.1093/nar/22.22.4673.

Vanathy, K., Parija, S.C., Mandal, J., Hamide, A., Krishnamurthy, S., 2017. Cryptosporidiosis: A mini review. Trop. Parasitol. 7, 72–80 https://doi.org/10.4103/tp.TP_25_17.

Wang, Y., Li, N., Guo, Y., Wang, L., Wang, R., Feng, Y., Xiao, L., 2018. Persistent occurrence of Cryptosporidium hominis and Giardia duodenalis subtypes in a Welfare Institute. Front. Microbiol. 9, 2830. https://doi.org/10.3389/fmicb.2018.02830.

Xiao, L., 2010. Molecular epidemiology of cryptosporidiosis: an update. Exp. Parasitol. 124, 80–89. https://doi.org/10.1016/j.exppara.2009.03.018.

Xiao, L., Escalante, L., Yang, C., Sulaiman, I., Escalante, A.A., Montali, R.J., Fayer, R., Lal, A.A., 1999. Phylogenetic analysis of Cryptosporidium parasites based on the small-subunit rRNA gene locus. Appl. Environ. Microbiol. 65, 1578–1583. https://doi.org/10.1128/AEM.65.4.1578-1583.1999.

Xu, Z., Guo, Y., Roellig, D.M., Feng, Y., Xiao, L., 2019. Comparative analysis reveals conservation in genome organization among intestinal Cryptosporidium species and sequence divergence in potential secreted pathogenesis determinants among major human-infecting species. BMC Genomics 20, 1–15. https://doi.org/10.1186/s12864-019-5788-9.

